## Supplementary material for "Preprinting the COVID-19 pandemic": Supp Model and Supp Tables 2 & 3

**Supplementary Model**

To investigate whether length of COVID-19 preprints compared to non-COVID-19 preprints was influenced by other factors, we constructed mixed-effects regression models. Two separate models were constructed examining a) all bioRxiv preprints with available word count data (n = 24159) and b) bioRxiv preprints that were published with available word count and publishing delay data (n = 3975). Word count was modelled as the outcome, and in all cases, model covariates included number of authors, preprint type (COVID-19 vs non-COVID-19) and date of posting (as calendar day of the year) specified as fixed effects, and country of corresponding author and bioRxiv category as random effects. Model a) contained additional fixed effect terms of a binary variable indicating whether the preprint was published or not, and an interaction between this variable and date of posting to adjust for increased likelihood of publishing over time since posted. Model b) contained additional fixed effect terms of publishing delay, and an interaction between this variable and preprint type, in order to investigate differential word count ~ publishing delay relationships for COVID-19 and non-COVID-19 preprints.

Significance of fixed effects were assessed via Wald tests and random effects were assessed via likelihood ratio tests (LRTs). All mixed regression models were conducted using function lmer() in R package ‘lme4’, v1.1-23 [1]. Confidence intervals were calculated as profile confidence intervals. Model collinearity was checked for by ensuring all variance inflation factors (VIF) were < 5 using function vif() in R package ‘car’, v3.0-10 [2].

Among all bioRxiv preprints, word counts were positively associated with authorship team sizes with an average increase of 72 words/author (Supplemental Table 2). There was significant evidence for a published:posting day interaction, suggesting that as the year went on, those preprints that were unpublished increased in length but those preprints that were published decreased slightly in length, though effect sizes were small (average increases of 1.4 words/day and decreases of 0.2 words/day, respectively). Word count also significantly varied with country of corresponding author and bioRxiv category, with intraclass correlations indicating these explained 3.5 and 4.2% of residual variance, respectively (Table S1). Adjusted for all these other factors, COVID-19 preprints remained significantly shorter than non-COVID-19 preprints, with an average difference of approximately 1600 words.

Limiting to those published bioRxiv preprints, similar associations/variances were observed with number of authors, country of corresponding author and bioRxiv category (Supplemental Table 3). There was a positive relationship between word count and publishing delay for both preprint types, such that an additional 100 words was associated with an average additional delay of 64.9 days for non-COVID preprints, but only 16.5 days for COVID preprints.

**Supplementary Model References**

1.	Bates D, Mächler M, Bolker B, Walker S. Fitting linear mixed-effects models using lme4. J Stat Softw. 2015;67: 1–48.

2.	Fox J, Weisberg S. An R companion to applied regression. Third. Thousand Oaks CA: Sage; 2019. Available: https://socialsciences.mcmaster.ca/jfox/Books/Companion/

**Supplemental Table 2. Outputs from mixed-effects regression predicting word count using all bioRxiv preprints. CI = confidence interval, ICC = intraclass correlation.**

| *Fixed effects* | | | |
| --- | --- | --- | --- |
| **Covariate** | **Coefficient (95% CI)** | **t** | **p (Wald)** |
| (Intercept) | 4447.1 (4205.4, 4685.2) | - | - |
| No. authors | 71.6 (66.4, 76.8) | 27.2 | < 0.001 |
| Preprint type = COVID-19 | -1592.8 (-1712.3, -1473.5) | -26.2 | < 0.001 |
| Preprint published = yes | 147.7 (5.7, 289.8) | 2.0 | 0.042 |
| Calendar day of posting | 1.3 (0.9, 1.7) | 6.9 | < 0.001 |
| Preprint published = yes * calendar day of posting | -1.5 (-2.7, -0.3) | -2.5 | 0.013 |
| *Random effects* | | | |
| **Covariate** | **ICC** | **Chisq** | **p (LRT)** |
| Country of corresponding author | 0.035 | 902.7 | < 0.001 |
| bioRxiv category | 0.042 | 481.9 | < 0.001 |

**Supplemental Table 3. Outputs from mixed-effects regression predicting word count using only published bioRxiv preprints. CI = confidence interval, ICC = intraclass correlation.**

| *Fixed effects* | | | |
| --- | --- | --- | --- |
| **Covariate** | **Coefficient (95% CI)** | **t** | **p (Wald)** |
| (Intercept) | 4638.4 (4311.2, 4964.6) | - | - |
| No. authors | 68.9 (56.4, 81.4) | 10.8 | < 0.001 |
| Preprint type = COVID-19 | -2224.0 (-2658.6, -1789.8) | -10.0 | < 0.001 |
| Publishing delay (days) | 1.5 (0.3, 2.8) | 2.4 | 0.017 |
| Calendar day of posting | 0.8 (-0.5, 2.0) | 1.2 | 0.213 |
| Preprint type = COVID-19 * publishing delay (days) | 4.5 (0.03, 8.99) | 2.0 | 0.049 |
| *Random effects* | | | |
| **Covariate** | **ICC** | **Chisq** | **p (LRT)** |
| Country of corresponding author | 0.023 | 100.9 | < 0.001 |
| bioRxiv category | 0.034 | 59.7 | < 0.001 |
